# Augmenting cognitive load during split-belt walking increases the generalization of motor memories across walking contexts

**DOI:** 10.1101/470930

**Authors:** Dulce M. Mariscal, Pablo A. Iturralde, Gelsy Torres-Oviedo

## Abstract

Cognitive load plays a role on the movement recalibration induced by sensorimotor adaptation, but little is known about its impact on the generalization of movements from trained to untrained situations. We hypothesized that altering cognitive load by distracting subjects during sensorimotor adaptation would facilitate the generalization of recalibrated movements beyond the training condition. We reasoned that awareness of the novel condition inducing adaptation could be used to consciously contextualize movements to that particular situation. To test this hypothesis, young adults adapted their gait on a split-belt treadmill (moving their legs at different speeds) while they observed visual information that either distracted them or made them aware of the speed difference between their feet. We assessed the adaptation and aftereffects of spatial and temporal gait features known to adapt and generalize differently when walking on the treadmill or overground. We found similar adaptation and aftereffects on the treadmill across all groups. In contrast, both groups with altered cognitive load (i.e., distraction and awareness groups) generalized their movements from the treadmill to overground more than controls, who walked without altered cognitive load. Of note, this effect was only observed in temporal gait features, which are less susceptible to online motor adjustments, and were eliminated upon experiencing large errors by briefly removing the split perturbation during adaptation (i.e., catch trial). Taken together, increasing cognitive demands during sensorimotor adaptation facilitates the generalization of movement recalibration, but this cognitive-mediated effect cannot eliminate the specificity of actions due to context-specific errors.

**New and Noteworthy:** Little is known about how cognition affects the generalization of motor recalibration induced by sensorimotor adaptation paradigms. We showed that augmenting cognitive load during adaptation on a split-belt treadmill led to greater recalibration of movements without the training device. However, this effect was eliminated when unusual motor errors were experienced on the treadmill. Thus, cognition can influence the generalization of sensorimotor adaptation, but it cannot suppress the context-specificity originated by the errors that one feels.

## Introduction

Generalization of learning is defined as the ability to apply knowledge acquired in one situation to new experiences. For instance, a tennis player will likely generalize the motor learning acquired from playing tennis to other sports played with rackets. This motor ability is studied in sensorimotor adaptation by assessing the carryover of movements recalibrated in a novel environment to variations of the training task (Ingram et al. 2000; Krakauer et al. 2000; Reynolds and Bronstein 2004; Cothros et al. 2006; Reisman et al. 2009; Torres-Oviedo and Bastian 2010; Wang et al. 2011; Torres-Oviedo and Bastian 2012; Bédard and Song 2013; Kitago et al. 2013; Howard and Franklin 2015; Wang and Song 2017). Notably, it has been shown that arm movements recalibrated when reaching in one direction generalize to reaching in other postures (Shadmehr and Mussa-Lvaldi 1994) or directions (Donchin et al. 2003; Malfait and Ostry 2004; Howard and Franklin 2015; Wang and Song 2017). On the other hand, the generalization of sensorimotor recalibration to movements without the training device is more limited (Kluzik et al. 2008; Reisman et al. 2009; Torres-Oviedo and Bastian 2010, 2012). This is, for example, evidenced by the reduced adaptation effects (i.e., aftereffects) following split-belt walking when stepping overground (untrained situation) compared to when stepping on the treadmill (trained situation) (Reisman et al. 2009; Torres-Oviedo and Bastian 2010, 2012). We are particularly interested in identifying factors regulating the generalization of sensorimotor adaptation because its translational value. Namely, the repetition of gait recalibration through split-belt walking can lead to reductions of gait asymmetry post-stroke (Reisman et al. 2013; Lewek et al. 2018), but it is critical that these improvements carryover to daily life situations. Thus, we are interested in factors mediating the generalization of sensorimotor adaptation to harness them such that motor improvements observed in clinical populations from these tasks generalize to untrained circumstances.

Previous findings indicate that context-specific cues from sensory, motor, or cognitive information regulate the generalization of sensorimotor adaptation. For example, sensory information specific to the adaptation condition will determine its generalization to other situations (Krouchev and Kalaska 2003; Wada et al. 2003; Osu et al. 2004; Ahmed et al. 2008; Ingram et al. 2010; Torres-Oviedo and Bastian 2010; Addou et al. 2011; Wang et al. 2011; Hirashima and Nozaki 2012; Howard et al. 2013). Similarly, actions before (Howard et al. 2012, 2013; Howard and Franklin 2015), during (Torres-Oviedo and Bastian 2012), or after (Howard et al. 2015) experiencing the novel condition regulate the generalization of recalibrated movements. Interestingly, cognitive load by altering subjects’ attention to the adapted task can also modulate the generalization of movements in reaching (Bédard and Song 2013). Thus, cognitive processes can alter the generalization of sensorimotor adaptation in volitional actions, but it remains unknown if this effect is also observed in more automated behaviors such as locomotion.

The idea that cognition can alter the generalization of locomotor adaptation is plausible given growing evidence that cognitive processes have an effect on sensorimotor adaptation. For example, changes in cognitive load during sensorimotor adaptation alter motor adjustments from one trial to the next (Taylor and Thoroughman 2008), the magnitude of aftereffects (Redding et al. 1992), or the rates at which individuals adapt (Bock 2010; Im et al. 2015) and de-adapt their actions (Malone and Bastian 2010). Thus, cognitive processes can modulate the recalibration of movements induced by sensorimotor adaptation, suggesting that it may also change the generalization of adapted movements. We particularly tested the hypothesis that altering cognitive load by distracting individuals would increase the generalization of motor patterns across walking conditions (i.e., treadmill vs. overground), whereas explicit information about the novel environment would reduce it. We further tested if the potential effects of altered cognitive load would be maintained when subjects experienced unusual errors in the training environment, which has been shown to limit the generalization of sensorimotor adaptation in reaching (Kluzik et al. 2008) and walking (Torres-Oviedo and Bastian 2012).

## Methods

### Subjects

A group of young adults were tested to investigate the effect of cognitive load during split-belt walking on the generalization of locomotor adaptation (experiment 1: n=30, 19 females; mean age 25.43± 1.53 yrs.). We also investigated the extent to which the effect of cognition on generalization was sustained upon experiencing large errors induced by briefly removing the split condition on a subsequent experiment (experiment 2: n=30, 19 females; mean age 27.28±1.37 yrs.). The study was approved by the University of Pittsburgh Institutional Review Board and it is in accordance with the Declaration of Helsinki. All subjects gave informed consent prior testing.

### Locomotor Paradigm

#### General Protocol

All participants performed a gradual split-belt paradigm consisting of a baseline, adaptation, and post-adaptation epochs (Fig. 1A). In the baseline epoch, two baseline blocks were collected: one for overground and another for treadmill walking to measure subjects’ regular walking in these two contexts. During the overground baseline block, subjects walked along an 8-meter walkway for 10 minutes at a self-selected speed. During the treadmill baseline block, subjects walked on the treadmill when the belts moved at the same speed (i.e., tied condition) at 1.125 m/s for 300 strides. A stride was defined as the time between two consecutive heel strikes (i.e., foot landing) of the same leg. In the adaptation epoch, subjects’ walked on the split-belt treadmill while the speed difference between the feet was gradually introduced (Fig. 1C). This was done to reduce the saliency of the split perturbation in the distraction group and the same was done in all other groups for consistency purposes. The general speed profile is illustrated in Figure 1C. In total subjects walked for 1200 strides. First, both belts moved at 1.125m/s for 300 strides, then one belt started to gradually speed up and the other to slow down during 600 strides until they reached a 2:1 belt speed ratio (i.e., fast belt moving at 1.5m/s and the slow belt moving at 0.75 m/s). The dominant leg (i.e., self-reported leg to kick a ball) walked on the fast belt. Lastly, the 2:1 split-belt ratio was maintained for 300 strides. In the post-adaptation epoch subjects walked overground and on the treadmill to assess the generalization and washout of the split-belt pattern, respectively (Fig 1B). During the overground post-adaptation block, subjects walked on a walkway for 10 minutes at a self-selected speed. Importantly, subjects were transported to the beginning of the walkway in a wheel chair to ensure we could record the initial overground steps following adaptation. Finally, during the treadmill post-adaptation block, participants walked with the two belts moving at the same speed of 1.125 m/s for 600 strides (figure1A). The initial steps during this epoch were used to quantify the remaining aftereffects that were not washed out by overground walking, and hence the remaining motor memory specific to the treadmill context (i.e., washout in Figure 1B).

**Figure 1:**
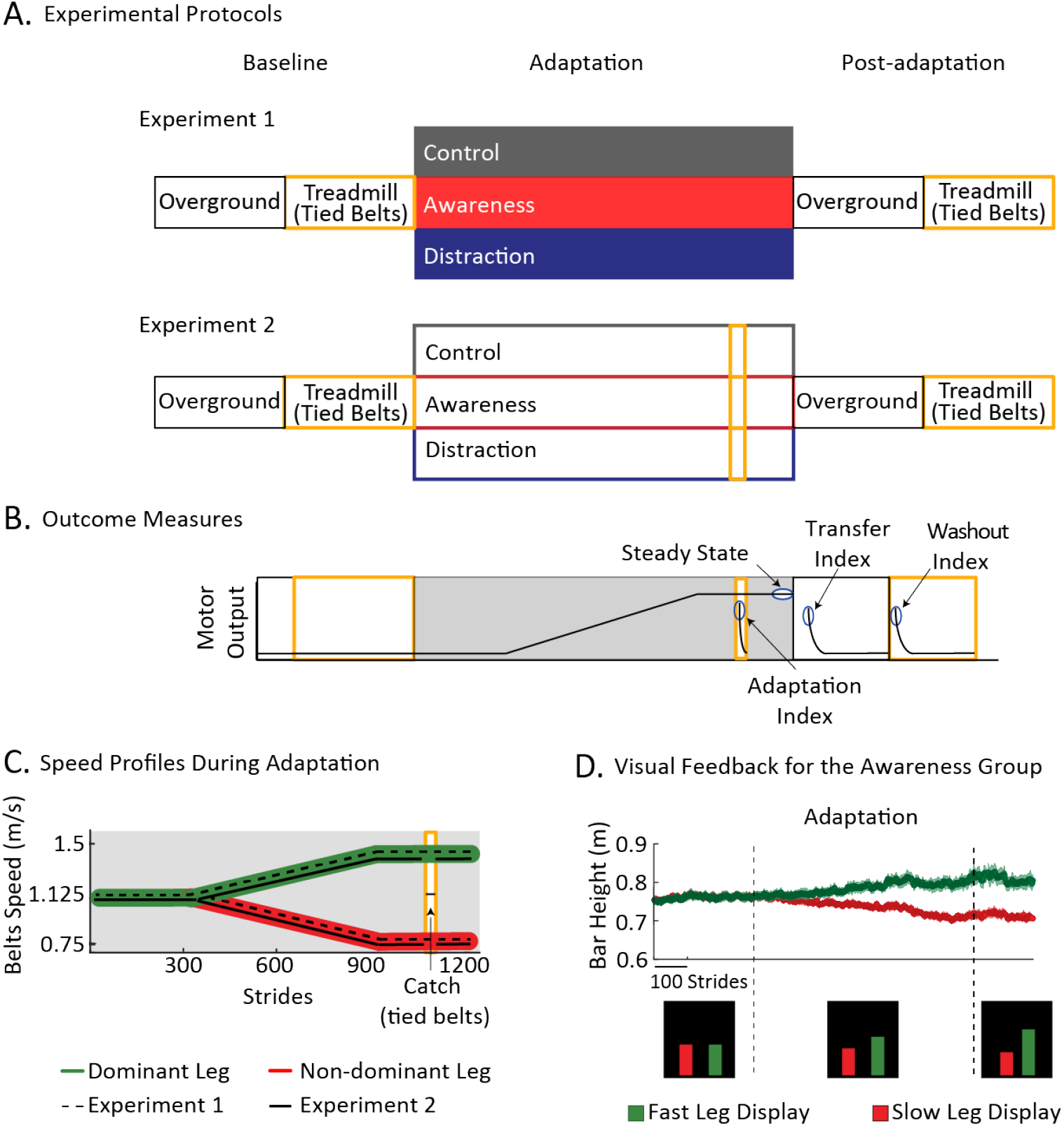
A. Experimental protocols. These consisted of three epochs: Baseline, Adaptation, and Post-adaptation, each of which had distinct blocks outlined with distinct colors. The adaptation block was further divided into three colors to indicate the distinct cognitive load experienced by each group: control (without altered cognitive load), awareness (with altered cognitive load by receiving information about speed difference between the feet) and distraction (with altered cognitive load by performing a secondary task unrelated to split-belt walking). Only the subjects tested in Experiment 2 experienced a 10-stride catch trial (two legs moving at the same speed) during the Adaptation epoch B. Outcome measures. Adaptation index, Steady State, Transfer index, and Washout index were collected at time periods indicated on of interest C. Speed profiles. We illustrate the time course of the speed at which the dominant (green) and non-dominant leg (red) walked during the Adaptation epoch. Speed profiles for the legs in Experiment 1 (black solid lines) and Experiment 2 (black dashed lines) are also presented to illustrate that only Experiment 2 had a catch trial during which both belts moved at 1.125m/s. D. Visual feedback that the Awareness groups received during Adaptation. Subjects observed progression bars that informed them about each foot speed. The averaged time courses ± standard errors are displayed for each bar. We also show snap shots of image that subjects observed during the pre-ramp, ramp, and hold phases during Adaptation.

For safety purposes, subjects held to a handrail during the very first few steps of the baseline, adaptation, and post-adaptation blocks on the treadmill until they felt comfortable walking with their arms unrestricted (as they walked during the overground blocks). Also, a plastic divider was placed between the treadmill belts to ensure subjects could not step on the wrong belt during treadmill blocks. Finally, all individuals wore a harness on the treadmill that only provided support in the event of a fall.

#### Experiment 1

To investigate the effect of cognitive load on the generalization of locomotor adaptation we tested three groups: distraction group (n=10), awareness group (n=10), and control group (n=10). The distraction and awareness groups were compared to the control group in which subjects adapted their gait without any instruction (Fig. 1A, gray). Subjects in the distraction group (Fig. 1A, blue) had altered cognitive load by performing a secondary task unrelated to split-belt walking. Specifically, they were required to count (with a handheld counter) the number of times that a specific word was mentioned in a TV show. We used this distraction procedure because it has been shown to have an impact on locomotor adaptation (Malone and Bastian 2010). On the other hand, subjects in the awareness group (Fig 1A, red) observed the evolution of the speed difference between their feet during the entire adaptation epoch. More specifically, these participants watched two vertical progression bars displayed on the left and right side of a screen placed in front of them (See snapshots of the screen on Fig 1D). This group was told that these bars corresponded to the speed of the left and right leg, respectively. Each bar’s height increased in real-time as the duration of the foot in contact with the ground increased. Figure 1D illustrates the time courses of the bars’ heights. These show that they were of the same height when the speed difference was zero and they were of distinct heights as the speed difference increased. We chose to display a biometric parameter instead of each belt’s speed because we wanted to use a measure that encompassed the walking speed variability. Further, we chose to display stance duration for each leg, rather than foot speed, because stance duration is a speed-related measure (Reisman et al. 2005)that could be displayed reliably, whereas foot speed was susceptible to marker occlusion. This visual feedback was created using a custom program coded with Vizard (Worldviz, Santa Barbara CA). Individuals were familiarized to the visual feedback with two short trials (~10 strides each) with the visual display while they walked at 1.5m/s and at 0.75m/s, which were the speeds for each foot in the full split condition. In addition, subjects also experienced a 300 pre-ramp phase with visual feedback and tied walking that served as a familiarization period before the two legs moved at different speeds. Lastly, individuals in the control and distraction groups wore a drape to prevent them from seeing their feet, whereas subjects in the awareness group did not. This was done to allow individuals in the awareness group to confirm the displayed speed difference by looking at their feet. All groups walked without visual stimuli during baseline and post-adaptation epochs.

#### Experiment 2

We ran a second experiment to investigate if the effect of cognitive load on the generalization of recalibrated movements was altered by large errors upon removal of the split condition. To this end, three additional groups were tested following the exact paradigm as in Experiment 1, but these groups also experienced large errors by briefly removing the split condition (i.e., catch trial) during the adaptation epoch. This catch trial was introduced 1050 strides into the adaptation epoch, so that there were 150 strides of walking at the 2:1 split ratio before and after this trial. It consisted of a 10-stride trial with the belts moving at the same speed (1.125 m/s, Fig 1C). The step length asymmetry (aftereffects) experienced during the catch trial were considered errors specific to the treadmill environment. Importantly, individuals were instructed to walk without holding the handrail on the treadmill during the catch trial, such that the steps on the treadmill and overground context were more comparable.

### Data collection

Kinematic and force data were recorded to characterize subjects’ gait. Kinematic data were recorded at 100 Hz with a Vicon Motion System (Oxford UK) and force data were recorded at 1000 Hz with an instrumented split-belt treadmill (Bertec, Columbus OH). Kinematic data was collected by measuring the position of reflective markers located bilaterally on the ankle (lateral malleolus) and hip (greater trochanter). Gaps in raw kinematic data due to marker occlusions were filled with a spline interpolation (Woltring; Vicon Nexus Software, Oxford Uk). Force data were used for detecting foot landing (i.e., heel-strike) and foot lifting (i.e., toe-off) in real-time to count strides and to determine the stance duration used in the visual feedback of the awareness groups. On the other hand, kinematic data were used to detect gait events on the treadmill and overground as in previous work (Torres-Oviedo and Bastian 2010, 2012). This was done such that the data analysis of these two walking contexts was more comparable given that we could not collect force data overground.

### Data Analysis

#### Gait Parameters

Step length asymmetry, known to robustly adapt during split-belt walking (e.g., Reisman et al. 2005), was used as a global measure to characterize gait adaptation and its generalization to overground walking. Step length asymmetry is defined as the difference of step lengths (anterior-posterior distance between ankle markers at heel strike) of two consecutive heel strikes and normalized by the sum of the step lengths (Eq.1). As a result, zero values represent symmetric step lengths, positive values indicate that the leg on the fast belt (i.e., fast leg) is taking longer steps than the slow leg, and vice versa for negative values.

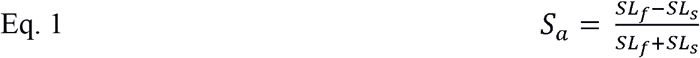

We also characterized spatial and temporal components of step length asymmetry (Eq. 2; Finley et al. 2015) because previous studies have shown distinct adaptation (Malone and Bastian 2010; Malone et al. 2012)and generalization (Torres-Oviedo and Bastian 2010; Sombric et al. 2017) of spatial and temporal gait features. Briefly explained, step length asymmetry can be decomposed into spatial (StepPosition, S_p_), temporal (StepTime, S_*t*_) and velocity (StepVelocity, *S_v_*) components of two consecutive steps (Eq. 2). StepPosition quantifies how far the foot lands away from the body when taking a step with one leg vs. the other (Eq. 3). StepTime compares the time to take a step (i.e., duration between two subsequent heel-strikes) with one leg vs. the other. This difference is scaled by the average velocity of the legs (Eq. 4). Lastly, StepVelocity quantifies the difference in speeds at which the foot moves with respect to the body when taking a step with one leg vs. the other. This difference is scaled by the averaged step time across the legs (Eq. 5).

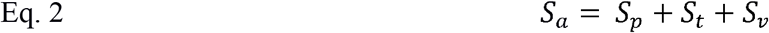

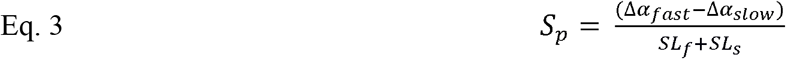

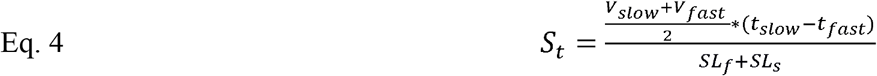

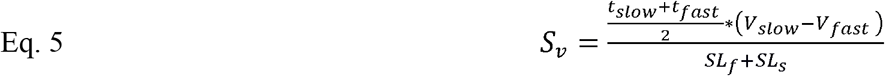

Where Δ*α_fast_* indicates the difference in distances between the fast leg’s landing position and the body at fast heel-strike and the previous slow leg’s landing position and the body at slow heel strike. Similarly, Δ*α_slow_* compares the distances between the slow leg’s landing position and the previous fast leg’s landing position (both with respect to the body location at slow and fast heel-strike, respectively). *t_slow_* quantified the duration between the fast leg’s heel-strike and the previous slow leg’s heel strike and *t_fast_* the duration between the slow leg’s heel-strike and the previous fast leg’s heel strike. Lastly, *V_fast_* and *v_slow_* represent the step velocity quantified as the relative velocity of the body with respect to the ankle in contact with the ground (i.e., fast ankle for *V_fast_* and slow ankle for *V_sow_*). Note that step length asymmetry and all its components are normalized by the sum of step lengths to account for differences in step sizes across individuals.

#### Outcome measures

Measures of subjects’ adaptation and generalization were computed for each of the gait parameters described above (i.e., Sa, Sp, St and Sv). Subjects’ adaptation performance was characterized with the steady state (SS) for each parameter and a global measure of extent of adaptation (AdaptExt). The steady state (*SS*) characterized subjects’ behavior at the end of the split-belt condition before they walked overground. This was computed using the average of the last 40 strides of adaptation (*Adapt_late_*) without the baseline bias (mean of last 40 strides of the treadmill baseline) as indicated in Eq. 6.

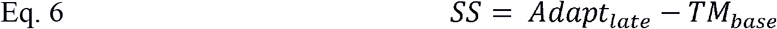

Extent of adaptation (*AdaptExt*) was used to measure the extent to which subjects counteracted the split-belt perturbation. This parameter was computed as the difference between the steady state for the step length asymmetry (*SS_a_*) and the steady state for the velocity component (*SS_V_*), which is a good proxy for the perturbation experienced by each subject (Finley et al. 2015). Formally expressed in eq 7.

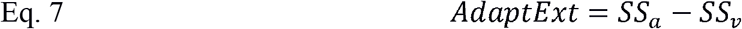

AdaptExt is always a positive measure since SSa monotonically increases from values neighboring SSv to zero values. Thus, large AdaptExt values indicated that subjects adapted their gait substantially on the split-belt condition, whereas small values indicated that they did not.

Aftereffects on the treadmill during the catch trial were used to compute an Adaptation index. These initial aftereffects were considered treadmill-specific errors since removing the split condition becomes a perturbation experienced on the treadmill after subjects experience a long adaptation period (Iturralde and Torres-Oviedo 2018) as in our protocol. Adaptation index was computed as the difference between the average of the first 3-strides during the catch trial (*TM_catch_*) and the treadmill baseline values (*TM_base_*).

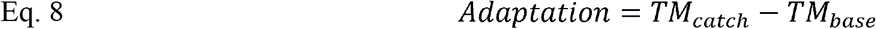

Generalization was characterized with two measures: 1) transfer index and 2) washout index. The transfer measure indicated the carryover of movements from the treadmill to the overground context. This measure was defined as the difference between the initial steps (averaged of the first 5-strides) overground after the adaptation epoch (*OG_after_*) and the baseline overground behavior (*OG_base_*) (Eq.9).

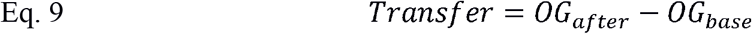

On the other hand, washout indicated the remaining aftereffects specific to the treadmill environment following overground walking. This outcome was quantified by the difference between the initial steps (first 5-strides) on the treadmill during the post-adaptation block (*TM_post_*) and the baseline behavior on the treadmill (*TM_base_*). Large values indicated little washout by the overground steps, whereas small values indicated substantial washout.

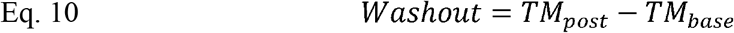

#### Statistical Analysis

We ran separate one-way ANOVAs on the distinct outcome measures to determine the effect of cognitive load alone (Experiment 1) and cognitive load in conjunction with large errors (Experiment 2) on the adaptation and generalization of gait. Fisher’s LSD post-hoc testing was used to compare the behavior across groups when we identified group main effects. We set the acceptable threshold for Type I errors to 5% in all statistical tests. Statistical analyses were performed with Stata (StataCorp, TX).

## Results

### Altered cognitive load did not affect the adaptation of gait

We observed that all subjects reached the same adapted state, regardless of their cognitive condition. This is qualitatively indicated by the time courses for all parameters during adaptation (Fig. 2A). Specifically, we did not find an effect of cognition load during the steady state in subjects who did not experience a catch trial (Steady State in Experiment 1 for Sa: F(2,27)=2.13, p=0.12; Sp: F(2,27)=1.15 p=0.33; St: F(2,27)=0.13 p=0.88) nor in those who did (Steady State in Experiment 2 for Sa: F(2,27)=0.73, p=0.49; Sp: F(2,27)=0.24, p=0.79; St: F(2,27)=0.91, p=0.41). These findings were further supported by the similar counteraction of the perturbation across groups with or without a catch trial (Experiment 1 without catch: AdaptExt F (2,27) =2.13, p=0.14 and Experiment 2 with catch: AdaptExt: F (2,27) =0.81, p=0.46, Fig. 2B). In sum, subjects’ cognitive state did not affect their ability to counteract gradual split-belt perturbations.

**Figure 2:**
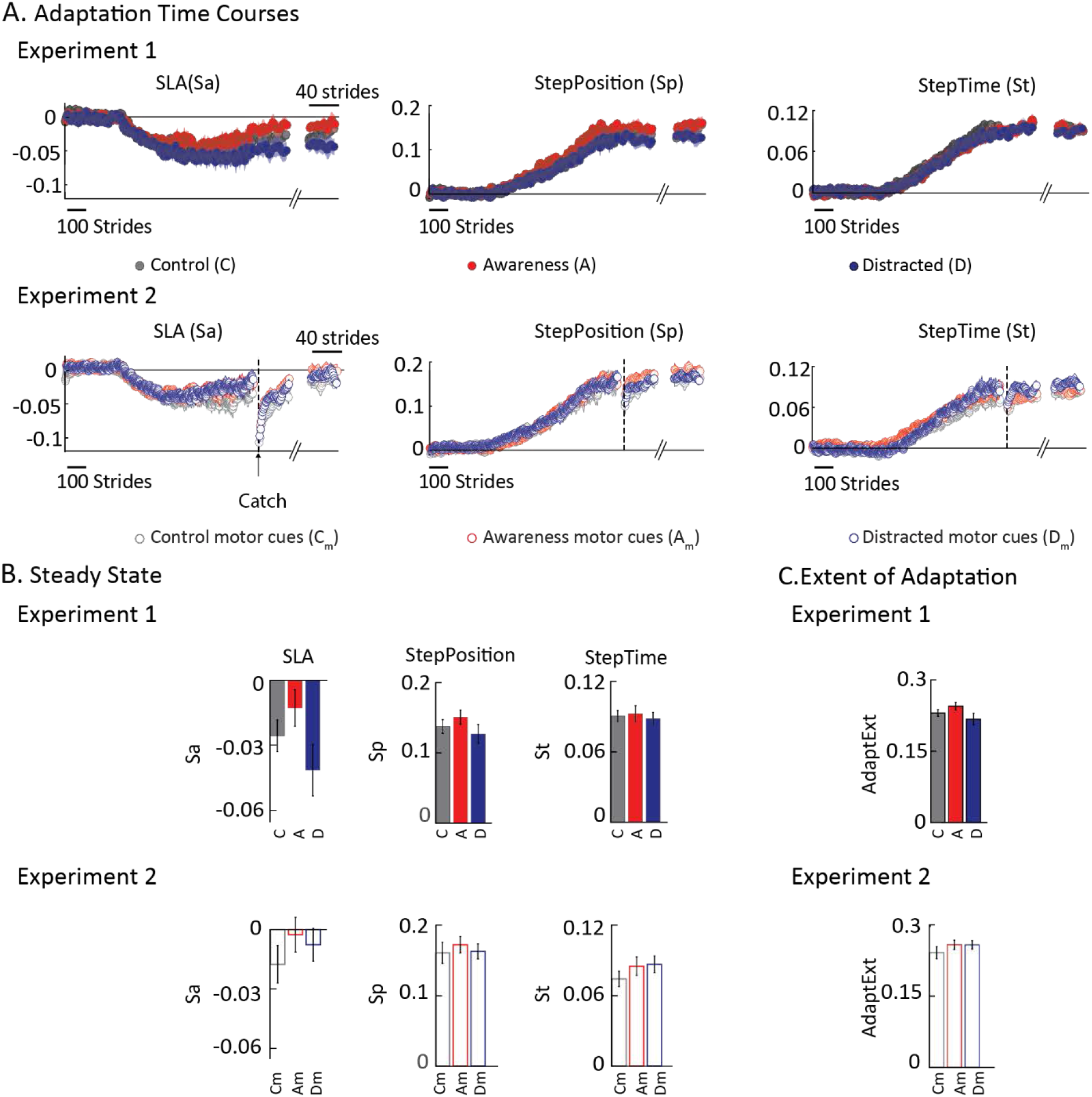
A. Time courses for each parameter during the Adaptation epochs of Experiment 1 (top row) and Experiment 2 (bottom row). B. Steady State at the end of the Adaptation epochs of Experiment 1 (top row) and Experiment 2 (bottom row). Bar plots indicate the mean adapted steady state per group ± standard errors. Note that we did not find a group effect for any gait parameter, indicating that cognitive load did not have an impact on the Steady State behavior prior to overground walking. C. Measure of adaptation extent for all groups. Bars’ height indicates the mean per group ± standard errors. All groups adapted their gait similarly.

### Altered cognitive load during split-belt walking increased the generalization of adapted step timing to overground walking

Cognitive load altered the aftereffects of step time overground. This is qualitatively shown by the distinct time courses of overground aftereffects in experiment 1 (Figure 3A). Note that the distraction and awareness curves (blue and red, respectively) have larger values than those in the control group (gray curve) for step time. This difference is also observed to a lesser extent in the time courses of step length asymmetry, but not of step position, for which curves overlapped across groups. Consistently, we found a significant effect of cognitive condition on overground aftereffects of step time (F(2,27)=5.51, p=0.01), but not for those of step length asymmetry (F(2,27)=1.55, p=0.23) or step position (F(2,27)=0.61, p=0.55). Further, post-hoc analysis indicated that the distraction and awareness groups had larger aftereffects overground in step time than the control group (control vs. distraction p=0.014, control vs. awareness p=0.005). Interestingly, there were no differences between the distraction and awareness groups (p=0.65). This suggests that the increased cognitive load in the awareness condition facilitated the generalization of motor adaptation, even if the secondary task provided contextual information about the treadmill. Taken together, we found that visual distractors during adaptation increased the transfer of updated step time on the treadmill to overground, but these differences in step time were not large enough to significantly change step length asymmetry overground.

**Figure 3:**
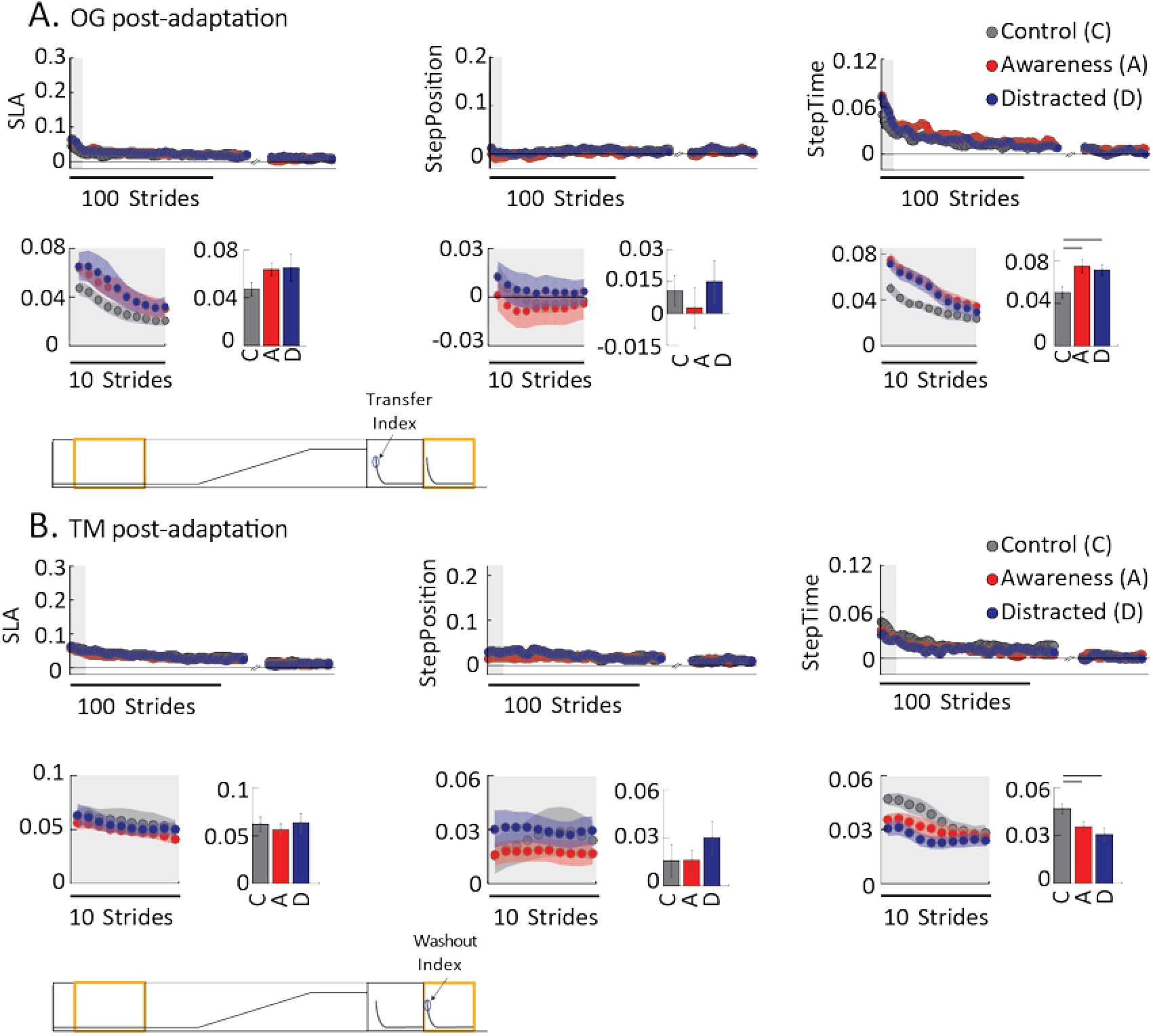
A. Stride-by-stride time courses and mean transfer values (i.e., overground aftereffects) are shown for all parameters during the post-adaptation block overground. B. Stride-by-stride time courses and mean washout values (i.e. remaining treadmill aftereffects) are shown for all parameters during the post-adaptation block on the treadmill. In both panels, gray shaded areas indicate the strides that are zoomed in the inserts. Each dot represents the average of 5 consecutive strides and colored shaded areas indicate the standard error for each group. Bar plots indicate either the mean transfer value (in Panel A) or the mean washout value (in Panel B) for each group ± standard errors. The black horizontal lines indicate significant statistical differences between groups. Recall that Experiment 1 was designed without a catch, thus aftereffects in the training context are not recorded for this group. For display purposes we use the axes are scaled as in Figure 4A presenting the aftereffects during catch for Experiment 2. This was done to qualitatively show that aftereffects overground and remaining aftereffects on the treadmill in Experiment 1 are much smaller than those observed during the catch.

The increased generalization of the adapted step time was further supported by the washout of treadmill aftereffects following overground walking. Figure 3B illustrates the time courses for subjects walking under different cognitive conditions. Note that groups adapted with altered cognitive loads during adaptation (i.e., distraction and awareness groups) showed smaller step time aftereffects when they returned to the treadmill, while their step length asymmetry and step position was similar across groups. Accordingly, we found a significant effect of cognitive condition on subjects’ remaining treadmill aftereffects following overground walking for step time (F(2,27)=5.47, p=0.01), but not for step length asymmetry (F(2,27)=0.22, p=0.81) or step position (F(2,27)=0.79, p=0.46). Moreover, post-hoc analysis on step time aftereffects indicated that subjects with altered cognitive load during adaptation had significantly smaller remaining aftereffects when they went back to the treadmill than those without it (control vs distraction p=0.003, control vs awareness p=0.034). This indicated that the distraction and awareness groups were more susceptible to washout from overground walking than controls. Once again, we did not observe differences between the distraction and awareness groups (p=0.33), further supporting that the increased cognitive load in the awareness group reduced, rather than facilitated, the context-specificity of locomotor adaptation, even if the secondary task provided explicit information about the unique split condition. Overall, our washout findings were consistent with our transfer results, in the sense that, the groups transferring the most were also those that had the least remaining aftereffects when returning to the treadmill. In sum, motor memories were more general when cognition was altered during adaptation not only because these memories carried over to an untrained situation, but because they were susceptible to walking in the untrained context (i.e., overground).

### Large errors during adaptation eliminated the effect of cognitive condition on generalization

The effect of cognition on the generalization of locomotor adaptation was not maintained when subjects experienced large errors induced by a catch trial during adaptation. This is indicated by the similar generalization and washout across cognitive conditions when experiencing a catch. We first noted that the cognitive condition did not have an effect on the treadmill aftereffects during the catch trial (Figure 4A. Sa: F(2,27)=2.24, p=0.13, Sp: F(2,27)=1.15, p=0.33, St: F(2,27)=0.87, p=0.43). These are the aftereffects that are experienced the very first time that the split condition is removed. Figure 4 also illustrates the time course of aftereffects when walking overground (Fig 4B) and when returning to the treadmill following overground walking (4C). Note that time courses for all groups overlap in all parameters and walking contexts. Consistently, there was not a significant effect of cognitive condition on overground aftereffects when subjects experienced a catch trial (Figure 4B. Sa: F(2,27)=0.36 p=0.70, Sp: F(2,27)=0.82, p=0.45, St: F(2,27)=0.95, p=0.4). Similarly, there was not a significant difference between the groups experiencing the catch trial on treadmill aftereffects following overground walking in all parameters (Figure 4C. Sa: F (2,27)= 0.39, p=0.68), Sp: F(2,27)=1.32 p=0.28; St: F(2,27)=1.61, p=0.22). Thus, all cognitive conditions had similar transfer and washout of treadmill aftereffects when they experienced large errors during adaptation.

**Figure 4:**
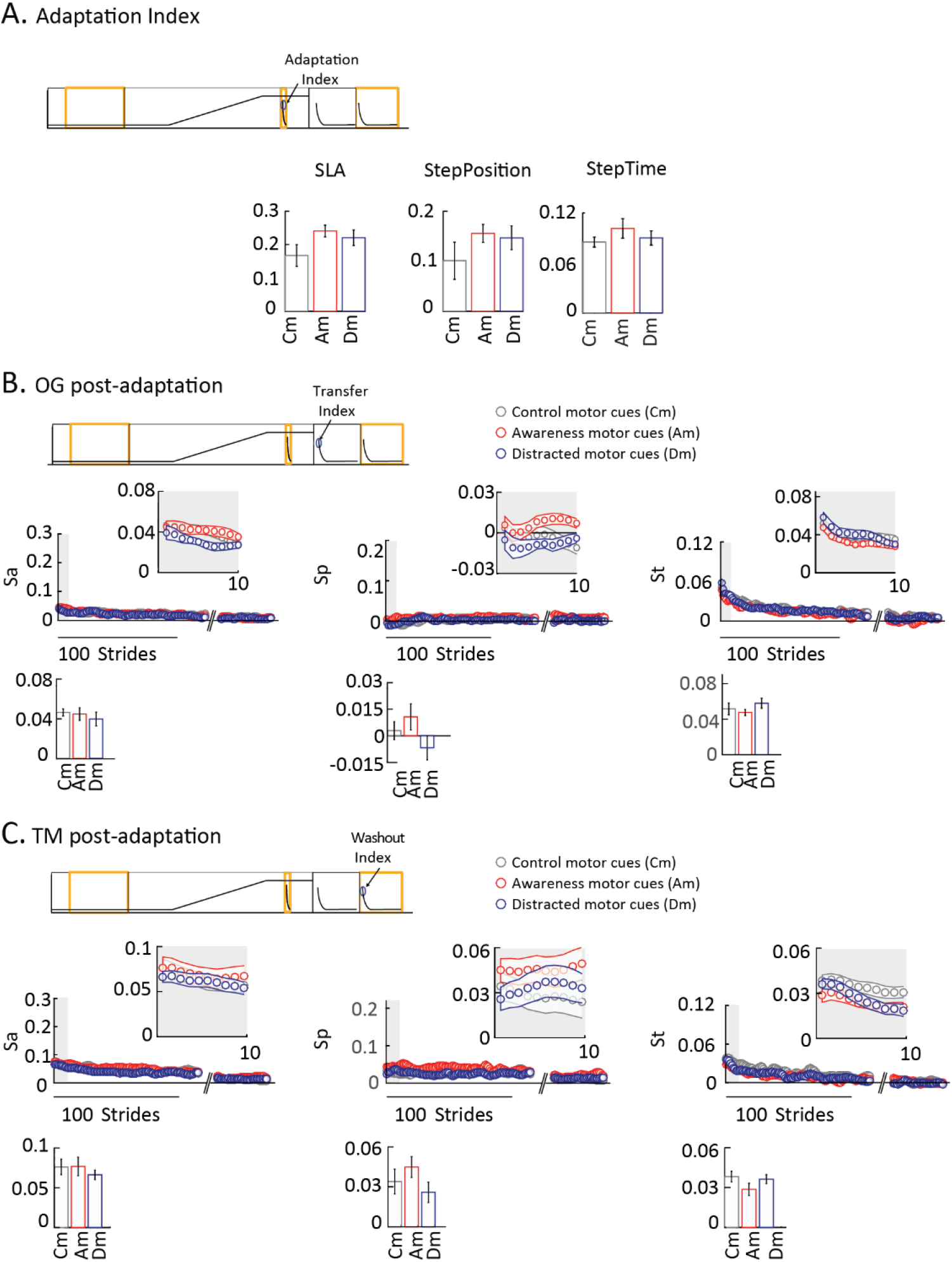
A. Mean adaptation index per group indicating the mean value for aftereffects experienced on the treadmill the first time that the split condition is removed. Error bars indicate standard errors. B. Stride-by-stride time courses and mean transfer values (i.e., overground aftereffects) are shown for all parameters during the post-adaptation block overground. C. Stride-by-stride time courses and mean washout values (i.e. remaining treadmill aftereffects) are shown for all parameters during the post-adaptation block on the treadmill. In both panels, gray shaded areas indicate the strides that are zoomed in the inserts. Each dot represents the average of 5 consecutive strides and colored shaded areas indicate the standard error for each group. Bar plots indicate either the mean transfer value (in Panel B) or the mean washout value (in Panel C) for each group ± standard errors. Cognitive condition did not have an effect on aftereffects on the treadmill and overground when large errors were experienced.

## Discussion

### Summary

We investigated how altering cognitive load during split-belt walking affects subjects’ ability to adapt and generalize gait movements. We also studied the effect of large errors during adaptation on the generalization of sensorimotor recalibration across different cognitive conditions. We found that cognitive load does not modulate subjects’ steady state in the split condition and the subsequent treadmill aftereffects. In contrast, cognitive condition had an impact on the generalization of temporal gait features adapted during split-belt walking. More specifically, augmenting the cognitive load during adaptation increased the generalization of aftereffects across walking contexts, even if the secondary task brought awareness to movements specific to the training condition. Interestingly, the effect of cognition on generalization was eliminated in the presence of large errors experienced during a catch trial (i.e., when the split condition was removed). Therefore, we find that a more general recalibration of walking occurs when cognitive resources during sensorimotor adaptation are occupied, but only in the absence of unusual errors in the training environment.

#### Cognitive load does not impact the sensorimotor adaptation to a gradual perturbation

We found that subjects’ performance during the adaptation epoch and subsequent aftereffects on the treadmill were not altered by cognitive load. These observations contrast previous findings indicating that increasing cognitive load limits subjects’ steady state performance (Ingram et al. 2000) or their ability to adjust movements from one trial to the next (Taylor and Thoroughman 2007, 2008). Altered cognitive load during locomotor adaptation has also been shown to slow down the adaptation rate (Malone and Bastian 2010). We believe that these differences stem from the distinct adaptation schedules in our study compared to previous work. More explicitly, our participants experienced a gradual perturbation, whereas the referenced studies were done in response to abrupt perturbations. Recent work indicates that cognitive-driven strategies, such as re-aiming contribute to motor performance upon large abrupt perturbations (Bond and Taylor 2015; Morehead et al. 2015). Perhaps we find that subjects’ performance to gradual perturbations is not susceptible to cognitive load because motor adaptation in this case requires less cognitive-based strategies.

Our results also showed that treadmill aftereffects, as measured in the catch trial, are not affected by the altered cognitive load. This is consistent with other walking studies (Malone and Bastian 2010; Long et al. 2016; Roemmich et al. 2016), but not with reaching literature showing that aftereffects are reduced when subjects perform cognitive task (Keisler and Shadmehr 2010). This discrepancy between reaching and walking could be explained by either 1) distinct contributions of explicit strategies to the adaptation of reaching and walking, or 2) distinct approaches for measuring aftereffects between these motor behaviors. First of all, consider that cognitive load likely influences aftereffects linked to explicit (i.e., strategic) corrections during adaptation, which may play a larger role in reaching than walking because reaching is a more volitional action. Second, aftereffects in walking are measured by removing the split perturbation (a.k.a., null condition), whereas aftereffects in reaching are measured by constraining the arm (a.k.a., error-clamp condition) (Keisler and Shadmehr 2010). As a result, feedback-mediated responses dominate aftereffects in walking (Iturralde and Torres-Oviedo 2018), but not in reaching. These feedback-mediated responses to unexpected transitions between walking conditions are more independent from cognitive processes than strategic actions (Malone and Bastian 2010). Therefore, cognition may only alter the explicit component contributing to aftereffects, but not the feedback-mediated one dominating aftereffects in walking.

#### Cognitive load during adaptation facilitates the generalization of motor adaptation

We found that increasing cognitive load during split-belt walking facilitates the generalization of adapted step timing, even when the secondary task brings awareness to movements specific to the training context. This was indicated by an increment on the generalization of step timing adapted on the treadmill and larger washout of this adapted step timing by overground walking; both of which observed in the distraction and awareness groups with increased cognitive load during adaptation. Our findings are consistent with inter-limb transfer literature showing that cognitive load during visuomotor rotations modulates the generalization of adapted reaches from one arm to the other (Kasuga and Nozaki 2011)and that explicit knowledge about the perturbation during adaptation does not disrupt generalization (Wang et al. 2011). We believe distractors might result in more generalized motor memories for two potential reasons. First, distractors might alter what is learned. We hypothesize that cognitive load reduces the explicit component of motor adaptation, which is tied to the environment, relative to the implicit one, which is tied to subjects’ actions and can be applied to other contexts. Second, large cognitive load might shift the credit assignment of errors during adaptation from the environment to oneself because subjects are more variable when cognitive resources are occupied. This potential change in credit assignment has been shown to alter the generalization of sensorimotor recalibration (Berniker and Kording 2008; Fercho et al. 2014). In sum, augmenting cognitive load during adaptation increases the generalization of learned movement across contexts because large cognitive load might alter the encoding of adaptation tied to subjects’ actions, rather than explicit corrections associated with the training environment.

We also found that cognitive load did not modulate the generalization of adapted step position. This observation is consistent with prior work showing that spatial and temporal aspects of gait generalize differently, and that the generalization of temporal gait features are easier to manipulate (Torres-Oviedo and Bastian 2010). This could be explained by the fact that during overground walking subjects could see their feet and these overrides aftereffects of step position, but not step timing. Notably, it has been shown that subjects use visual information to adjust their foot placement when taking a step, but not step timing (Marigold et al. 2008; Matthis and Fajen 2014; Maeda et al. 2016). Thus, we might not observe the influence of cognitive load on the generalization of step position because of the reliance on online feedback control for foot placement when walking overground.

#### Large errors increase the context-specificity of locomotor patterns

We observed that large errors upon removing the split condition override the impact of cognition on aftereffects. This was shown by the similar aftereffects between groups experiencing large errors during a catch trial, regardless of whether subjects walked overground or on the treadmill. These results are consistent with previous work showing that large errors during adaptation limit the generalization of aftereffects when walking overground (Torres-Oviedo and Bastian 2012). It has also been shown that subjects can switch faster between locomotor patterns when they experience transitions from split to tied walking (Malone et al. 2011; Sombric et al. 2017; Day et al. 2018). Therefore, aftereffects overground and on the treadmill might be reduced because errors during the catch trial might facilitate the transitioning between split and regular walking patterns.

#### Clinical implications

Our results might have an impact on the rehabilitation of hemiparetic gait because error-augmentation protocols, like the one presented here, can induce gait improvements in stroke survivors (Reisman et al. 2007; Savin et al. 2014) that persist with repeated exposure (Reisman et al. 2013; Lewek et al. 2018). However, if treadmills and robots are to be used for correcting patients’ movements, it is critical that the learned movements carry over to “real-life” situations beyond the training context. Here, we show that increasing cognitive load during sensorimotor adaptation facilitates the generalization of adapted behavior to different environments. These findings are promising for two reasons. First, individuals with motor disorders are often trained by either bringing self-awareness to their motions and explicit instructions on how to move (Lewek et al. 2018). Our results suggest that the generalization of motor improvements from these motions with large cognitive load will not be limited. Second, our results suggest that sensorimotor adaptation protocols, like split-belt walking, might lead to more general motor improvements if patients adapt their movements with increased cognitive load. However, future work is needed to test this hypothesis. In conclusion, our results suggest that increased cognitive load during rehabilitation therapies might lead to encoding more general motor memories, whereas errors specific to the training environment tied them to the training situation.

## Acknowledgments

Supported by NSF 1342183, NSF 1535036

